# SHAMAN: bin-free randomization, normalization and screening of Hi-C matrices

**DOI:** 10.1101/187203

**Authors:** Netta Mendelson Cohen, Pedro Olivares-Chauvet, Yaniv Lubling, Yael Baran, Aviezer Lifshitz, Michael Hoichman, Amos Tanay

**Affiliations:** Department of Computer Science and Applied Mathematics Weizmann Institute of Science, Rehovot 76100 Israel

## Abstract

Genome wide chromosome conformation capture (Hi-C) is used to interrogate contact frequencies among genomic elements at multiple scales and intensities, ranging from high frequency interactions among proximal regulatory elements, through specific long-range loops between insulator binding sites and up to rare and transient cis‐ and trans-chromosomal contacts. Visualization and statistical analysis of Hi-C data is made difficult by the extreme variation in the background frequencies of chromosomal contacts between elements at short and long genomic distances. Here we introduce SHAMAN for performing Hi-C analysis at dynamic scales, without predefined resolution, and while minimizing biases over very large datasets. Algorithmically, we devise a Markov Chain Monte Carlo-like procedure for randomizing contact matrices such that coverage and contact distance distributions are preserved. We combine this strategy with bin-free assessment of contact enrichment using a K-nearest neighbor approach. We show how to use the new method for visualizing contact hotspots and for quantifying differential contacts in matching Hi-C maps. We demonstrate how contact preferences among regulatory elements, including promoters, enhancers and insulators can be assessed with minimal bias by comparing pooled empirical and randomized matrices. Full support for our methods is available in a new software package that is freely available.

## INTRODUCTION

In order for chromosomal conformations to be robustly studied using the Hi-C^1,2^ technology, a large pool of nuclei is fixed, digested and chromosomal proximity events are recovered through massive sequencing of ligated restriction fragment ends. The data is then summarized as a contact matrix, in which the number of recovered ligations between every pair of restriction fragments is recorded. Since the total number of potential restriction fragment pairs in the genome is very large (at the order of hundreds of trillions), and since most restriction fragments contact with extremely low frequencies, any Hi-C matrix, including those sequenced at the maximal depth reported so far (billions of reads)^34,5^, is very sparse. Subsequent Hi-C analysis is thereby statistical, pooling together contacts across ranges of restriction fragments, or *bins*, to ensure data is transformed into statistically robust features (e.g., contacts between bins). Following binning, additional statistical modeling is usually performed to distinguish specific contacts of pairs of loci or regions from the background distribution of contacts over chromosomes, or to support informative visualization of the data.

Modeling chromosomal contact background distribution^6^ can be approached parametrically, by assigning bias-parameters to fragment-ends and modeling the contact probability given such parameters^7,8,9,10^. Alternatively, a non-parametric approach considering the marginal coverage for genomic bins can be applied to normalize systematic biases that are linked with one-dimensional sequence‐ or epigenetic‐ preferences^11,12^. Modeling chromosomal domain structure and identifying insulation hotspots can be approached based on such normalized binned matrices^3,5^ or performed by comparison of empirical contact distributions to a parametric model^4^. Searching for contact hotspots can similarly be performed by comparing the expected number of contacts given a parametric model^4,13,14^, by adapting techniques for peak finding^3,15,16^ on appropriately binned matrices, or by pooling contacts over specific pairs of epigenomic features^17,13^. Current methodologies for computational analysis of Hi-C matrices have allowed identification of key chromosomal features, including topological domains and CTCF/cohesin loops, and are becoming more sensitive as Hi-C matrices become richer in contacts, and there is a large number of software packages and algorithms^18,19,20^, most of which are not mentioned here for lack for space, suggesting different computational strategies for detecting these features in large matrices.

Despite much progress in the computational processing of Hi-C maps, their universal modeling remains challenging, in particular when searching for differential contacts between maps or when combining analysis of short range and long-range contacts. For example, the parametric models that were proposed require strong assumptions on the factors affecting contact distributions, and can be computationally demanding, while the a-parametric approaches that are based on binning, limit the resolution of the assay, and require careful additional normalization or stratification of chromosomal distance effects and other factors. In particular, a universal approach for comparing empirical features of Hi-C contact maps to those expected given simple assumptions is lacking.

In this paper we develop an approach for shuffling Hi-C contacts to generate randomized matrices that conserve the empirical number of contacts per restriction fragment and the empirical distribution of genomic distances over contacts. We then introduce techniques for analyzing Hi-C matrices by comparing observed and randomized data, based on an easy to use K-nearest neighbor approach for assessing differential contact densities. Our randomization algorithm and analysis suite is available through a new R package (called *shaman*), and we demonstrate its use by reanalyzing Hi-C data on mouse embryonic stem cells and human cancer cell lines, deriving high resolution models for contact insulation, loops, and longer range contacts.

## Results

### Shuffling Hi-C matrices while preserving contact marginal and distance distributions

The contact probability between pairs of restriction elements in Hi-C matrices is strongly correlated with the inverse genomic distance between them. When analyzing this trend globally (**Fig 1a**), ranges of genomic distances that grow exponentially (1-2kb, 2-4kb, 4-8kb,…, 64-128MB, 128-256MB) are populated by a comparable number of Hi-C contacts on average despite varying in size by five orders of magnitude. More precisely, logarithmic distance bins show a fivefold relative enrichment over this trend for the distance range between 100kb and 1MB, an effect strongly coupled to the topological domain structure of chromosomes^4,5,21,22^. The genomic distance effect on Hi-C contact probability, and its variation in specific distance ranges, is likely to reflect the true physical dynamics of the chromosome fiber, but it poses a difficult analytic challenge since the contact frequencies represented within Hi-C matrices can vary over a factor of 10,000 and more. A second major factor affecting the distribution of contacts in a Hi-C matrix is the overall efficiency of the complete protocol (fixation, digestion, ligation, retrieval, sequencing and mapping) for each restriction fragment, which can be estimated by the total number of contacts per fragment, denoted here as the *marginal coverage* (**Fig 1b-c**). This compound effect can span a range of over 100 fold, which can be correlated with the fragment sequence composition, chromosomal accessibility, digestion efficiency, overall fragment length, and sequencing mapability^7^. Differences in marginal coverage can increase the number of contacts between favored fragments in a way that is indirectly correlated with various epigenetic factors, thus introducing complex technical biases in the data.

**Figure 1:**
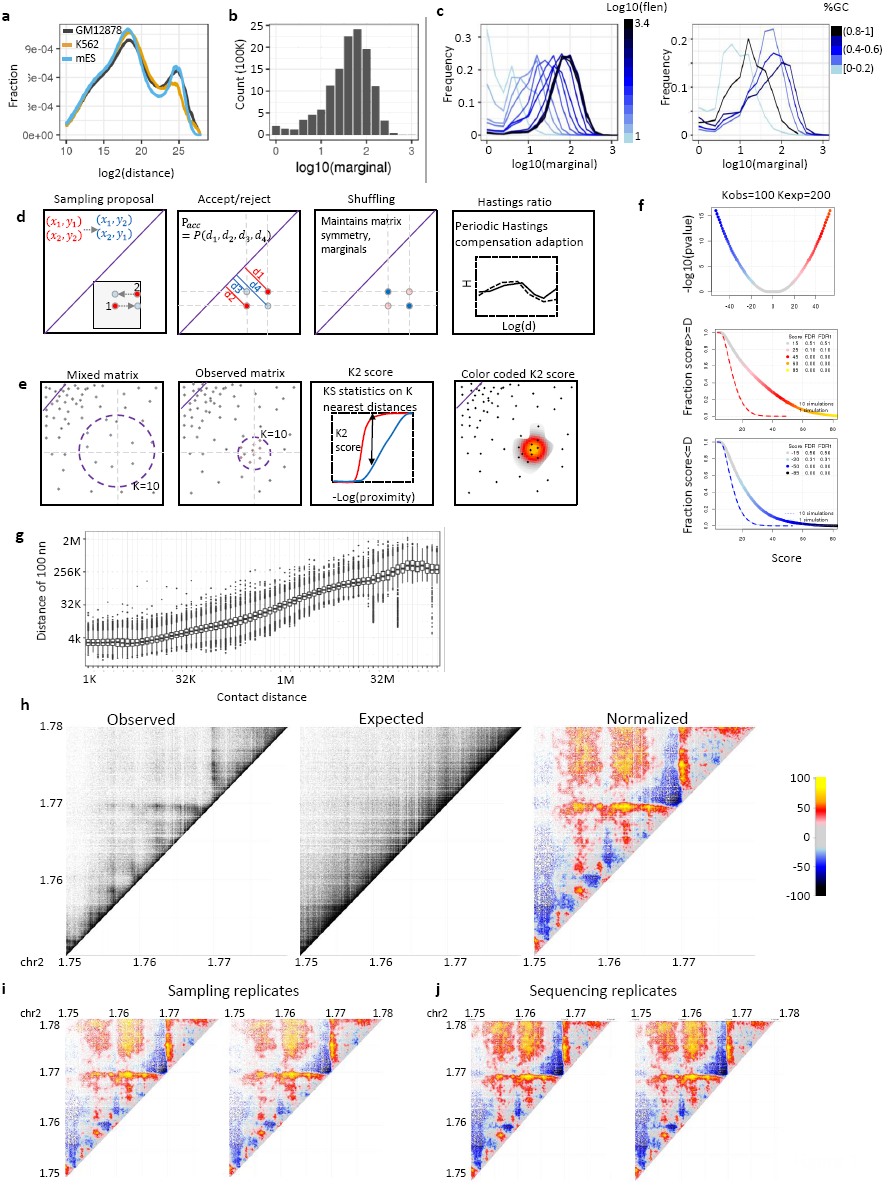
Bin-free multiscale HiC D normalization. **a) Decay curve** ‐ probability of contact as a function of log linear genomic distance, shown for Rao et al human GM12878 and K562 and Olivares et al mouse ES Hi-C. **b)** Distribution of marginal coverage, number of contacts per fragment, for Rao et al K562 Hi-C. **c)** Similar to b, stratified by fragment length (left) and by fragment G/C content (right). **d)** MCMC-like randomization algorithm scheme. Refer to methods for detailed description. In short, a pair of near-by contacts are randomly proposed for coordinate interchanging based on the probabilities of the distances between the contact end points **e)** D score computation. For each observed contact, a score is computed by comparing the K-nearest neighbor distances in the observed and shuffled matrices using Kolmogorov Smirnov D statistic. Refer to methods for detailed description. **f)** D score significance. Kolmogorov Smirnov (KS) D statistic color coded by value for KS test with observed K=100 and shuffled K=200 values (top). FDR for positive scores (D>0) (middle) and negative scores (D<0) (bottom) as computed by fdr test (Methods). **g)** Distribution of log distance to 100^th^ nearest neighbor stratified by log linear distance computed for K562 Hi-C data. **h)** Shown are observed (left), randomized (middle) and color coded D score normalized maps for the K562 HOXD region. **i)** Replicate sampling of contacts in the K562 HOXD locus, generating a randomized matrix and computing the D normalized score. **j)** Replicate randomization of contacts in the K562 HOXD locus, followed by D score computation.

Randomizing a Hi-C matrix, so as to preserve both genomic distance and marginal contact distributions cannot be achieved by naïve shuffling of contact pairs (which can preserve marginal contacts density but not the genomic distance distribution), or by sampling from a parametric model assuming the empirical contact distance distribution (which will not be consistent with the marginal coverage). We therefore developed a Markov Chain Monte Carlo (or Metropolis-Hastings-like) randomization algorithm, by repeatedly sampling pairs of contacts and shuffling them by swapping the fragment end points, with a ratio proportional to their contact distance probabilities, correcting for the asymmetric distribution of sampled and shuffled distances using an adaptive procedure (**Fig 1d**, Methods) (**Fig S1a**). The resulting shuffled matrix, after symmetrizing all contacts, maintains twice the precise marginal coverage for each restriction fragment, as well as nearly exactly the same distribution of contact distances observed in the empirical contact map (**Fig S1b**). We used sampling of pairs with restricted total genomic distance (methods) to improve overall sampler efficiency (**Fig S1c**). Parallelizing shuffling over chromosomes allows complete randomization of a Hi-C experiment with 1 billion reads within 7 hours on a standard machine with a core for each shuffled chromosome. Importantly, such randomization is performed once per dataset as an analysis pre-process following sequence mapping and contact calling. Applying our algorithm to dense Hi-C matrices generated for K562 and GM12787 human cancer cell lines^3^, and to a Hi-C matrix generated from mouse ES nuclei^23^, confirmed that randomized matrices closely follow the empirical contact distance distributions (**Fig S1b**)

### Using D score for bin-free comparison of observed and randomized contact densities

To assess relative enrichment of contacts around a point of interest in the Hi-C matrix, we define a Euclidean proximity metric between contacts (**Fig 1e**) and compute for each observed contact, the cumulative distributions of proximities over the K contact pairs closest to the point of interest in the empirical and randomized matrix. This can be performed efficiently even for very large matrices, using standard algorithms for identifying the K nearest neighbors in a metric space. We defined the *D* score of each observed contact in the HI-C matrix as the Kolmogorov-Smirnov (KS) D statistic obtained by comparing the empirical and randomized distributions of distances.

We note that the background distribution of the D scores depends on the selection of K, where larger K implies more statistical power but lower resolution. We determine statistical significance of D values using reshuffling and computing of False Discovery Rate (FDR) values (**Fig 1f**).

This algorithmic approach supports analysis at very high resolution for short-range contacts with high density where the K nearest neighbors are observed within a very short range of the contact of interest (**Fig 1g**, **S1d**). The same strategy uses much lower effective resolution for very long-range contacts with low density, and we are guaranteed natural scaling of the resolution as sequencing coverage increases. To visualize contact densities we simply color code points on the Hi-C matrix according to their D scores (**Fig 1h**), with a color scale that is selected according to FDR scores to ensure reproducibility and robustness (**Fig 1i-j**). In summary, the combination of matrix randomization and bin-free analysis of contact density allows flexible visualization and statistical analysis of complex Hi-C matrices, which naturally scales with the chromosomal distance of assayed contacts.

### Comparing Hi-C matrices

One of the main challenges of analyzing Hi-C data is comparing contact maps acquired in different conditions or from distinct cell types, and distinguishing between the technical differences affecting captured contacts distributions and true differential contact enrichment with possible regulatory impact. Using the randomization approach described above, we implemented two strategies for comparing contact matrices. First, we can use the D scoring scheme for comparison of two empirical matrices instead of relying on it for correcting one empirical matrix with a randomized one. When using this approach, difference in marginal coverage between the maps, or differences in the overall contact distance distribution will affect, and may even dominate the differential signal. Second, we can correct each dataset independently for effects associated with marginal coverage and contact distance distribution and compare the resulted D scores following normalization. This is exemplified by analysis of contact distributions around the beta globin locus in erythroid K562 cell line and lymphoblastic cell line GM12878. As expected, the active conformation of this site in K562 (**Fig 2a**) is markedly different from the repressed GM12878 conformation (**Fig 2b**), with marked increase in the contact intensity on several loops closing up the globin genes and the locus control region. The direct comparative analysis (**Fig 2c**) highlights the gain of globin-associated loops (H1-H3) and a specific loop closing up the downstream domain (H4). The normalized comparison **(Fig 2d)** suggests that increased looping intensity in H2-H3 is less specific, while H1 and H4 remains highly specific even after correcting each map separately to contact distance and marginal distribution.

**Figure 2:**
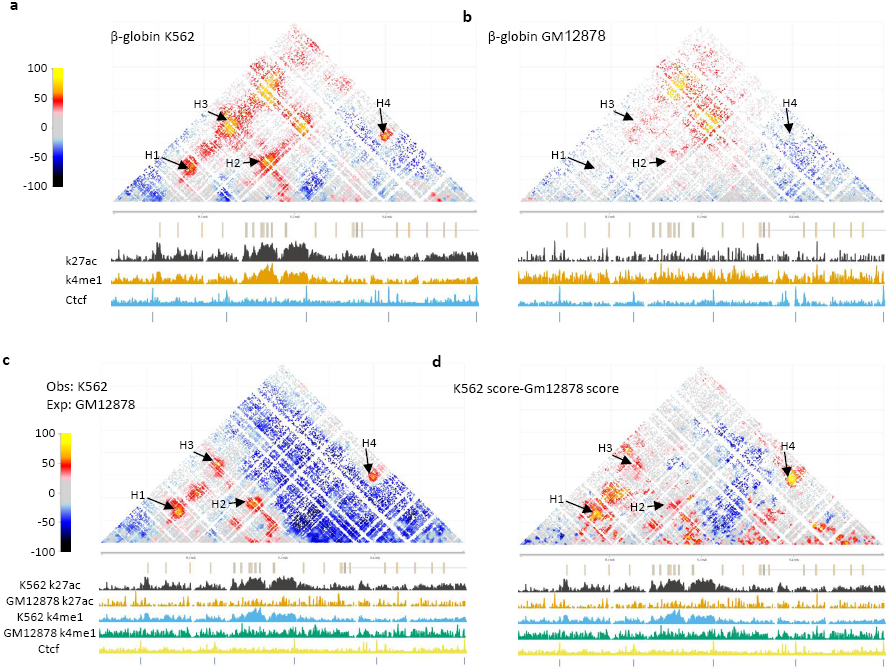
Comparing Hi-C matrices. **a)** Normalized D map of the K562 active beta-globin locus, annotated with genes, K562 H3K27ac and H3K4me1 profiles, as well as ES CTCF. **b)** Similar to a for GM12878. **c)** Similar to a, for D score map generate by comparing K562 observed Hi-C with GM12878 observed Hi-C. **d)** Similar to c, comparing K562 and GM12878 maps by subtracting normalized scores computed for each dataset separately (by comparing to its randomized matrix) at union of all points from both datasets.

Additional differential looping below the H4 loop is observed in the normalized comparison, suggesting direct comparison may mask short-range differential contact due to global changes in contact distribution (See **Fig S2** for similar analysis erythroid/megakaryocytic and lymphoid regulated loci). In summary, comparison between Hi-C maps is simplified through the D scores, but its implementation and interpretation, with or without normalization of the compared matrices, should be adapted to the analytic question at hand.

### Analysis of contact distributions around epigenetic landmarks

Pooling together Hi-C sub-matrices around pairs of epigenetic hotspots was used before to characterize the potential for preferential long-range chromosomal interaction in specific contexts, such as CTCF/cohesion sites^3,24^, transcription factor binding sites^17,25,26^ and polycomb repressive domains^27–29^. A challenge in this approach is to ensure proper normalization, since systematic biases for variable marginal coverage on epigenetic hotspots, or overall differences in chromosomal distance distributions between classes of such hotspots, may skew the pooled matrices considerably. This for example, frequently results in the appearance of enriched horizontal and central bands (a cross-like pattern), which represent a combination of marginal coverage on the examined hotspots, rather than specific enrichment in the pairing probability. Using randomized Hi-C matrices as statistical background allows for straightforward control over marginal and genomic-distance effects (**Fig 3a**, Methods). Ratio of pooled observed and randomized contacts around potentially contacting CTCF binding sites confirm the previously reported enrichment of contacts around CTCF sites with converging motifs (**Fig 3b**). Importantly, CTCF sites contact enrichment is observed nearly symmetrically around the focal contact point (**Fig 3c**), with significant enrichment observed at least 5kb from the motifs on either side (see spatial pattern for pairs filtered on significant contact enrichment in **Fig S3a**). The intensity of CTCF contact enrichment strongly depends on the genomic distance (**Fig 3d**), where 2-fold enrichment over the background is observed on average for sites at 100kb distance, but no enrichment (or in fact, mild negative enrichment) is observed for sites within 1MB distance, possibly due to closer competing CTCF pairings. Analysis of divergent sites (reverse-to-forward) shows nonspecific enrichment of contacts in the lower-right quadrant (representing contacts of genomic elements between the two divergent sites) suggesting that the occurrence of forward-to-reverse pairs between these sites combine together to generate indirect and non-specific contact enrichment pattern (**Fig S3b**). Pairs of CTCF sites with matching orientation (forward-to-forward, reverse-to-reverse, **Fig 3e**) are associated with contact enrichment on a band, which is likely to represent pairing of a hidden site, reverse oriented for a forward pair (left), forward oriented for a reverse pair (right), which give rise to a non-specific enrichment downstream from the first forward site or upstream from the second reverse site (right) (**Fig S3b**). This analysis demonstrates how indirect effects (due to convergent CTCF sites or other factors located near the analyzed loci) can give rise to seemingly significant contact enrichment. The specificity of factor interaction must therefore be supported by a localized, centered enrichment in spatial analysis, as observed for convergent CTCF sites here.

**Figure 3:**
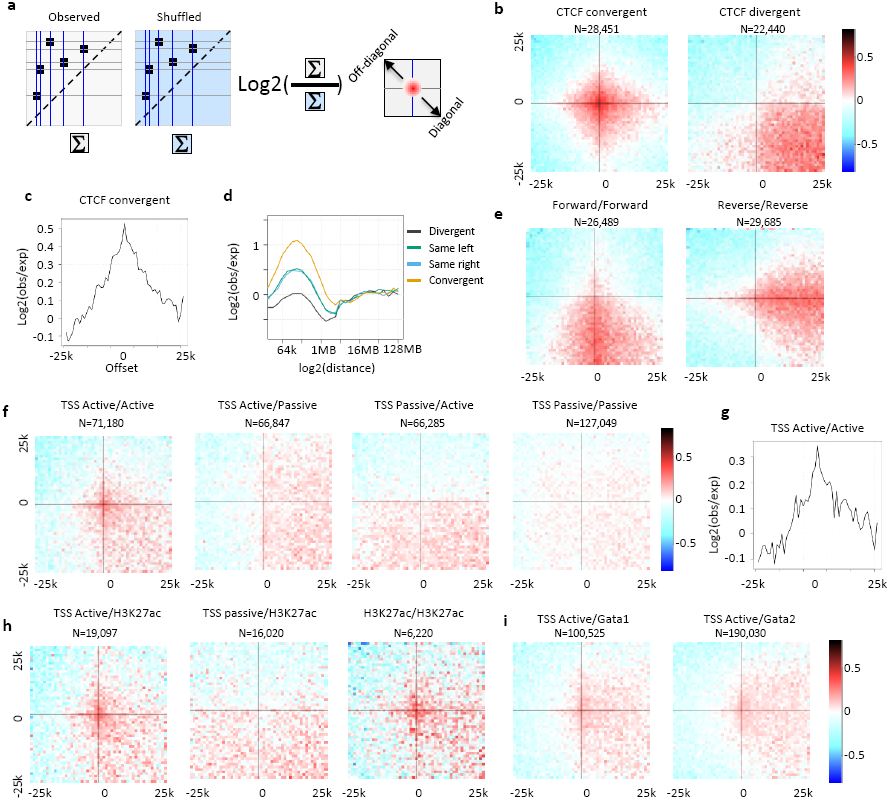
Contact distributions around epigenetic landmarks. **a)** Schematic overview for pooling of contacts around epigenetic hotspots and comparing observed to randomized distributions. **b)** Spatial enrichment ‐ shown are log2(obs/shuffled) number of contacts 25kb around CTCF convergent (top left), CTCF divergent (top right), CTCF forward/forward (bottom left) and CTCF reverse/reverse (bottom right) that are distanced between 200K and 2MB from each other and that contain a D score>30 anywhere within the 50K window at 1kb resolution (Methods). **c)** CTCF convergent enrichment profile (log2(obs/shuffled)), by offset from hotspot. **d)** Contact enrichment at CTCF pooled pairs stratified by log linear distance. **e)** Similar to b for TSS. **f)** Similar to b for contact enrichment between active TSS**. g)** Similar to b for active TSS and H3K27ac (left), passive TSS and H3K27ac (middle), and H3K27ac and H3K27ac (right). **h)** Similar to b for active TSS and Gata1 (left) and Gata2 (right).

In addition to CTCF sites, it has been proposed that the three-dimensional folding of chromosomes can bring distant regulatory elements such as promoters and enhancers into close spatial proximity. Applying the statistical method described above to contacts above 200kb and within less than 2Mb associating a TSS with another genomic element (excluding CTCF sites, Methods), we observe a modest (1.41 fold) but significant (chi-square, p<<0.001) localized enrichment in contacts between active pairs of TSS (**Fig 3f-g**), or between active TSSs and putative H3K27ac marked enhancers (**Fig 3h**) and Gata1,2 binding sites (**Fig 3i**). Non-specific enrichment downstream active TSSs (i.e. transcribed gene bodies) vs. passive TSSs may represent a general tendency of such region to contact at longer range (**Fig 3f**, middle panels). Pairing of H3K27me3 or H3K9me3 hotspots in K562 or GM12878 were not associated with significant local enrichment (**Fig S3c**). In conclusion, spatial analysis of pooled putative interaction is a powerful and sensitive tool for distinguishing truly synergistic and specific pairing from various background effects, and can be used to critically assess hypothesis on short‐ and long-range regulatory contacts and hubs.

### Screening for long-range contacts using D scores

Using the D scoring scheme, it is possible to screen for contact enrichment hotspots throughout the entire Hi-C matrix without prior assumptions. We applied this approach to the high depth lymphoblasts and ertirhroid maps, deriving 462 and 1304 non-overlapping contact enrichment hotspots, respectively ranging in distance from 4Mb to 100Mb (Methods). We observe that many of the identified hotspots were characterized by a broad pattern of enrichment, associating together genomic elements at a much larger scale than observed above for CTCFs or TSSs. To quantify this observation, we extracted for each contact hotspot a 2D matrix reflecting the spatial contact enrichment pattern within a 0.5MB range at 50kb resolution. We then clustered these matrices and studied the average spatial pattern associated with each cluster (**Fig S4**). This confirmed the broad nature of nearly all long-range contact hotspots we identified, showing contact enrichment at distance range of 100kb around the center, and in many of the cases larger distances are observed. Highly localized contacts between elements genomically separated by more than 4MB are very rare, and are all linked with possible assembly or mapping limitations (**Fig S4c**). These results suggest that weak preferential association between TADs and compartments, time of replication effects, cell cycle phase effects or other global phenomenon, all of which are not corrected by our randomization, may underlie a significant fraction to the very long range contact enrichment we observed. Spatial analysis as performed above is essential in these cases, in order to ensure observed contact enrichment between certain local elements is not a mere consequence of their embedding into broadly enriched contacting domains.

## DISCUSSION

We introduce *Shaman* ‐ a new approach for analyzing Hi-C contact matrices based on a matrix randomization algorithm combined with a bin-free scheme for analysis of contact density enrichment. *Shaman* is using its randomization scheme for balancing the Hi-C challenging background contact probability distribution, providing effective multi-scale visualization of the data so as to capture CTCF-mediated looping, promoter-enhancer contacts, topological domain boundaries and long range, low-specificity contacts between entire topological domains, all within one scheme and without any parameter setting. Randomizing matrices that preserve simultaneously the empirical marginal and contact distance distribution is also facilitating statistical analysis at various regulatory contexts, as effects at any epigenomic context or any mixture of chromosomal proximities can be studied systematically by comparison of empirical and randomized contact densities.

The resolution of Hi-C screen depends on sequencing depth and the genomic distance spanned between elements whose interaction is being screened. As shown in Fig 1g, the typical range of the k-nearest contacts to a point of interest increases with distance and thus making detection of effects that are localized but enriched only mildly above the background level increasingly difficult or impossible. Targeted methods, such as 4C^9,30,31,25^, ChiA-PET (REFs) or Capture Hi-C^32–34^ can enrich for specific contacts and increase resolution in loci of interest. Nevertheless, detection of putative interaction based on targeted methods can be confounded when applied to long range loci, since enrichment of contacts between two loci can be a consequence of regional and non-specific effects. Multiple sources of indirect effects, including replication time and cell cycle biases, variable compartment and pseudo-compartment preferences^22^ or changes in the overall intensity of enriched contacts between active loci may result in detection of statistically significant contacts that are nevertheless non-specific. We suggest that unbiased Hi-C analysis using the tools introduced here can provide a global map of such global effects, and highlight regions for potential follow up by targeted methods, which can then be interpreted not only locally, but given the broader context. Moreover, when assessing overall preferences of contacts among families of loci (e.g. transcription factor binding sites), it is essential to distinguish specific vs. non-specific contact enrichment by controlling foci of putative interaction with spatial analysis (as demonstrated in Figure 3, and while controlling for overall capture preferences of assayed elements). The data here suggest that Hi-C maps with reasonable sequencing burden should be generated routinely prior to application of targeted approaches, and that a combined non-specific and focused study design can achieve maximum impact and ensure the interpretability of the derived putative interactions.

## ACKNOWLEDGMENTS

Research was supported by the European Research Council (EVOEPIC), and by the NIH 4DN program.

## Software availability

The SHAMAN package is available from https://tanaylab.bitbucket.io/shaman.

## METHODS

### Estimating the contact genomic distance probability distirubtion

Focusing on one chromosome at a time, and given a set of Hi-C intra-chromosomal contacts between coordinates (x_i_,y_i_) we estimate the contact distance decay curve D by binning contacts using floor(B*log2(|x_i_-y_i_|), B is set by default to:

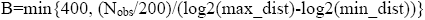

Where N_obs_ is the total number of contacts and *min_dist*, *max_dist* define the range of genomic distances observed. For each bin, we count the number of observed contacts to generate the raw decay curve *D_i_*. We smooth the decay curve by defining a smoothing window of size S bins in each direction, such that the smoothed value *DSmooth* for bin S < I < max_bin-S is defined:

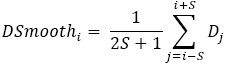

Setting S= min{20, (B/10)} provides good tradeoff between curve stability and its resolution. We apply linear regression to compute the smoothed S end bins on each side, taking into account 2*S observed raw bins, and correcting for negative values by setting to 0.

### Randomizing Hi-C matrices using a Metropolis-Hastings scheme

Given an observed Hi-C matrix for a given chromosome, we define a probability distribution over all symmetric matrices that preserve the observed total number of contacts per column/row by considering only the contact chromosomal distances:

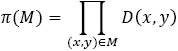

We implement a Metropolis-Hastings (MH)-like algorithm to sample from this probability distribution. In general, the sampler selects (or “proposes”) contact pairs at random (e.g., (a,b), (c,d)) and tries to cross them (e.g., to (a,d),(c,b)) with probability that is proportional to their chromosomal distance probabilities. Sampling arbitrary contact pairs would typically generate low probability (high distance) crosses, so we implemented a more restricted pair sampling scheme, by selecting the first contact (a,b) uniformly from all current matrix contacts, and the paired contact uniformly among all pairs (c,d) such that |a-c|<W and |b-d|)<W. Here W is defined as the proposal distribution *grid size*. Such sampling was implemented efficiently by binning contacts into two dimensional bins of size W and repeatedly sampling a paired contact from bins overlapping or adjacent to (a,b)’s bin until a contact within distance W was sampled.

In a simple Metropolis algorithm, the proposal function is symmetric, hence once we sample a proposed change to the matrix, accepting the change with probability min(1,π(M)/π(M’)) guarantees convergence to the desired distribution. However when the proposal distribution Q(M→M’) is non symmetric, as in our case, one must correct for the asymmetry by multiplying the acceptance probability by the Hastings ratio h(M,M’) = Q(M’→M) / Q(M→M). The non-symmetry in our proposal distribution stems from the sampling of contact pairs from a matrix that is highly imbalanced in its contact distance distribution, generating longer-distance crosses with higher probability. Computing the hastings ratio explicitly is possible, if we can count at each sample the ratio between total number of contacts around the original and proposed (crossed) contacts, but this may be computationally expensive. Given the definition of our probability distribution, we choose to approximate the Hastings factor *h* using only the two contacts (a,b),(c,d) selected for shuffling (rather than the entire matrix M), and to further parameterize our approximation, using the chromosomal distances involved d_1_=|a-b|, d_2_=|c-d|. d_3_=|a-d|, d_4_=|c-b|, setting h ((a,b),(c,d)) = h_d_(d_3_)*h_d_(d_4_)/h_d_(d_1_)*h_d_(d_2_). Here h_d_ is correcting the proposal distribution asymmetry by assuming it is defined completely by the sampled and shuffled contact distances. Importantly, the approximated Hastings effect as captured by h_d_ is imperfect, and we must periodically update it to avoid continuous drift of the sampled matrices.

Our sampling algorithm therefore works as follows. We assume the original contact matrix M has N_obs_ contacts following an estimated distance decay curve D:

1) Initialize h_d_. We do this by sampling a large number (0.5N_obs_) of contact pairs M and measuring the distance of all putative crossed contacts M_rand_(x) (without performing any shuffle). h_d_(x) is defined as the ratio M_rand_(x)/M, where the actual value is derived for 200 logarithmic bins and smoothed, as performed generically for estimating D.
2) Until we accept (0.0001 N_obs_) shuffles: sample contact pairs as described above and shuffle them with acceptance probability:

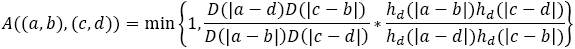
3) Correct the hastings ratio according to the accumulated skew of the current sampled matrix M from the expected distance decay curve. This is done by correcting for the divergence of the current randomized matrix curve (Dobs) from the original expected one:

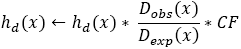 Where the default value for the correction factor CF= 0.25.
4) We go back to step 2 and repeat iteration until K_burn_*N_obs_ shuffles are accepted, where K_burn_ is the burn-in parameter that we typically set to 40 for a proposal grid size of 500K followed by another 40 iterations for proposal grid size of 1Mb.
5) After randomization is complete, we symmetrize the resulted Hi-C matrix, such that any contact (a,b) is duplicated to (b,a). The final matrix thereby contains 2N_obs_ contacts.

### Characterizing Hi-C contact density enrichment using the D score

We assess the density of a Hi-C matrix at a point (a,b) by characterizing the sequence of Euclidean distances from (a,b) to the K contacts most proximal to it. This can be computed efficiently using a K-nn data structure. The distance sequence, or the cumulative distribution defined by it can be easily compared between matrices using a Kolmogorov Smirnov (KS) D statistic, provided the K parameter is scaled according to the total number of contacts in the matrix, hence setting 2K neighbors for the randomized matrix. We define this D statistic, when comparing observed and randomized matrix, as the density enrichment score, where positive D values indicate contact enrichment over the expected background distribution and negative D values represent depletion. The parameter K can be tuned according to the overall coverage, where smaller K values can enhance resolution but may decrease statistical power.

To determine the background distribution of D values we randomized a matrix and compared its D score over all contacts against re-randomized matrices. This distribution was used as a background in order to assess FDR values (Fig 1f).

### Epigenetics landmarks

We downloaded epigenetic data from ENCODE as described in Table S1. For each epigenetic landmark, we screened for genomic regions with strong ChIP-seq signal (peak percentile as listed in table S1).

- For CTCF, we annotated each peak according to the motif strength, requiring motif strength > 99.988% of the genome-wide distribution. Each 20bp window of the CTCF peaks was classified as C (only ChIP), F (ChIP + forward motif), R (chip + reverse motif) or B (ChIP + forward and reverse motifs). Only F and B CTCF were considered for contact distributions analysis of CTCFs.
- For TSS, H3K4me3 peaks that were within 5K of transcription start site and were at least 5k from CTCF chip peak were marked as active TSS. TSS that were not marked with H3K4me3 (distance to H3K4me3 peak > 5k) or CTCF (distance to CTCF chip peak>5k) were considered as passive TSS.
- For all other epigenetic ChIP-seq, peaks were filtered for CTCF peaks (distance > 5k).

### Screening contact hotposts

For a given pair of epigenetic landmarks (e.g. CTCF_forward, CTCF_reverse), we define hotspot candidates as all possible pairs distanced 200Kb-2Mb from each other. Candidates were further filtered by contact enrichment in the normalized contact map, requiring a D score higher than 30 within a 50x50 Kb window centered on the hotspot. Spatial enrichment is computed by pooling all sub-matrices around candidates at 500 bp resolution in both observed and random data and computing the log ratio for each spatial bin at 1k resolution.

### Clustering contacts hotspots spatial structures (fig S4)

We screened for all long-range contacts (distanced between 4-100M) with high score (>80). High scoring contacts were merged if they shared a K-nearest-neighbor (K=100), defining hotspots, and the center of each hotspots was defined. For each hostspot, we defined a grid of 20x20 100kb bins around the center and extracted the maximum D score in each bin to create a vector of features characterizing the spatial structure around the hotspot. We then clustered hostpots using Kmeans, and visualized the derived patterns as shown in Fig S4.

---

**Table S1:** Epigenetic landmarks from ENCODE.

| Cell type | ChIP-Seq | From | Peak percentile |
| --- | --- | --- | --- |
| K562 | CTCF | ENCODE/UW CTCF Binding | 0.999 |
| K562 | H3K27ac | ENCODE/Broad Histone | 0.999 |
| K562 | H3K4me1 | ENCODE/Broad Histone | 0.999 |
| K562 | H3K4me3 | ENCODE/Broad Histone | 0.995 |
| K562 | Gata1 | ENCODE/SYDH TFBS | 0.995 |
| K562 | Gata2 | ENCODE/SYDH TFBS | 0.995 |
| K562 | H3K9me3 | ENCODE/Broad Histone | 0.999 |
| H1 | H3K27me3 | ENCODE/Broad Histone | 0.999 |

**Supplementary Figure 1.**
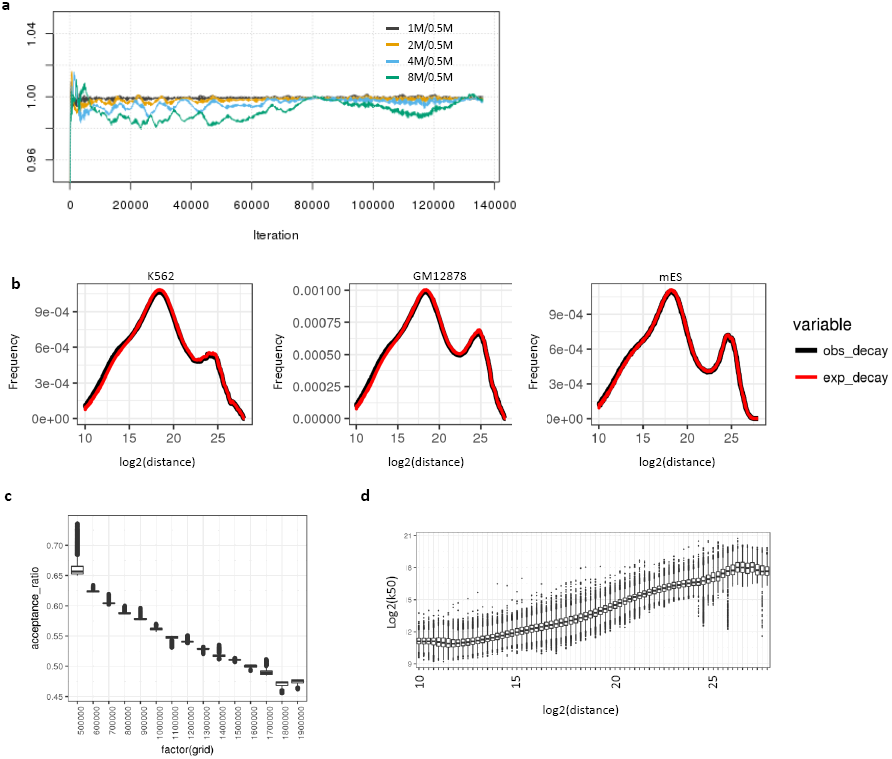
**a)** Hastings ratio per correction for several color coded log2 distance bins. **b)** Decay curve for observed (black) and randomized (red) datasets, K562 (left), GM12878 (center), mouse ES (right). **c)** Monte Carlo Markov Chain acceptance probability distribution as a function of the distance W between the proposed contact points (Methods). **d)** Distribution of log distance to 50^th^ nearest neighbor stratified by log linear distance computed for K562 Hi-C data.

**Supplementary Figure 2.**
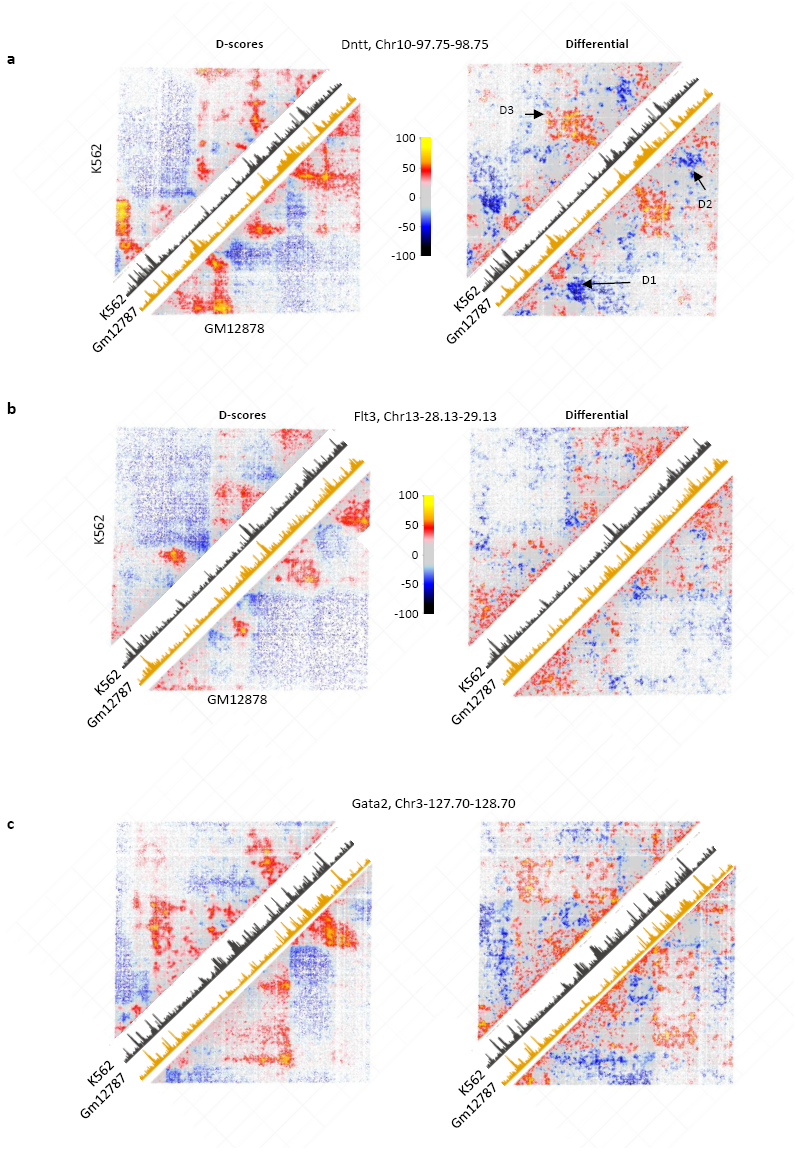
**a-c)** Direct comparison of D scores maps is shown to the left, K562 data is top left, GM12878 is bottom-right. K27ac ChIP-seq data is shown for the two cell types on the diagonal. Visualizing the differential D score (as discussed in Figure 2) is shown on the right, (symmetric matrix is shown, where enrichment (ref-yellow) represent denser contacts in K562 and anti-enrichment (blue) represent denser contacts in GM12878.

**Supplementary Figure 3.**
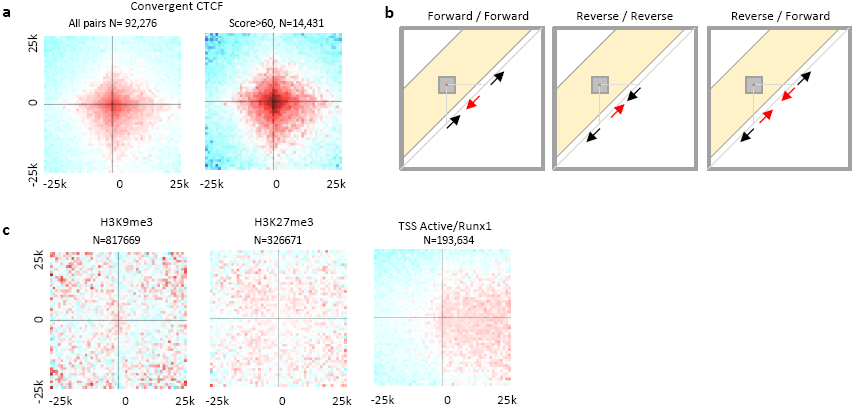
**a)** Shown are log2(obs/shuffled) number of contacts 25kb around any CTCF convergent sites (left) and CTCF convergent sites that contain D score>60 anywhere within the 50K window that are distanced between 200K and 2MB from each other at 1kb resolution (Methods). **b)** Schematic visualization explaining the spatial contact enrichment patterns of non-convergent CTCF pairs, including forward-to-forward (left), reverse-to-reverse (center) and reverse-to-forward (right), all rely on convergent CTCFs to generate the observed spatial pattern. **c)** Similar to a for H3K9me3 sites (left) and H3K27me3 sites (center) and active-TSS and Runx1 (right) requiring D score > 30 anywhere within the 50K window.

**Supplementary Figure 4.**
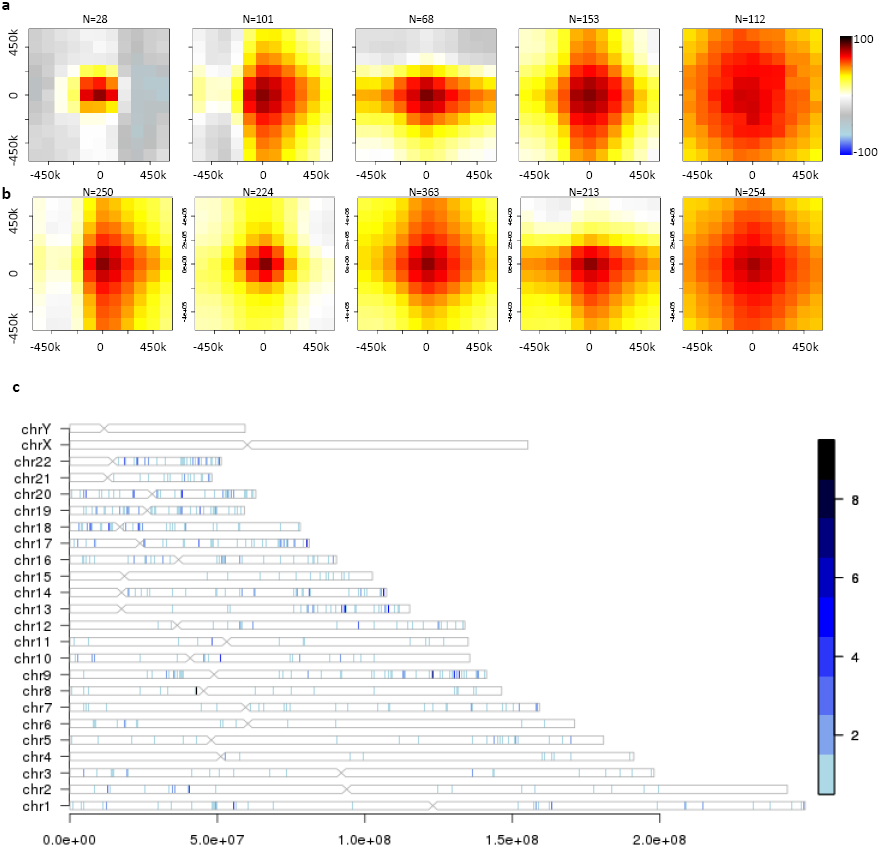
**a)** Long range contact enrichment hotspots clusters average spatial profile in K562. **b)** similar to a for GM12878 dataset. **c)** Chromosomal ideogram depicting all genomic loci with D score > 80.

